# Global non-random abundance of short tandem repeats in rodents and primates

**DOI:** 10.1101/2021.11.24.469825

**Authors:** Masoud Arabfard, Mahmood Salesi, Yazdan Hassani Nourian, Iman Arabipour, Ali Mohammad Ali Maddi, Kaveh Kavousi, Mina Ohadi

## Abstract

**Background:** While of predominant abundance across vertebrate genomes and significant biological implications, the relevance of short tandem repeat (STR) abundance to speciation remains largely elusive and attributed to random coincidence for the most part. In a model study, here we collected whole-genome abundance of mono-, di-, and trinucleotide STRs in nine species, encompassing rodents and primates, including rat, mouse, olive baboon, gelada, macaque, gorilla, chimpanzee, bonobo, and human. The obtained unnormalized and normalized data were used to analyze hierarchical clustering of the STR abundances in the selected species.

**Results:** We found massive differential abundances between the rodent and primate orders. In addition, while numerous STRs had random abundance across the nine selected species, the global abundance conformed to three consistent <clusters>, as follows: <rat, mouse>, <gelada, macaque, olive baboon>, <gorilla, chimpanzee, bonobo, human>, which coincided with the phylogenetic distances of the selected species (p< 4E-05). Exceptionally, in the trinucleotide STR compartment, human was significantly distant from all other species.

**Conclusion:** We propose that the global abundance of STRs is non-random in rodents and primates, and probably had a determining impact on the speciation of the two orders. We also propose the STRs and STR lengths which specifically coincided with the phylogeny of the selected species.

## Introduction

With an exceptionally high degree of polymorphism and plasticity, short tandem repeats (STRs) (also known as microsatellites/simple sequence repeats) are a spectacular source of variation required for speciation and evolution[1–3]. The impact of STRs on speciation is supported by their various functional implications in gene expression, alternative splicing, and translation[3–10].

STRs are a source of rapid and continuous morphological evolution[11], for example, in the evolution of facial length in mammals[12]. These highly evolving genetic elements may also be ideal responsive elements to fluctuating selective pressures. A role in evolutionary selection and adaptation is consistent with deep evolutionary conservation of some STRs, as “tuning knobs”, including several in genes with neurological and neurodevelopmental function[13].

While a limited number of studies indicate that purifying selection and drift can shape the structure of STRs at the inter- and intra-species levels [14–19], the global abundance of STRs at the crossroads of speciation remains largely unknown.

Mononucleotide and dinucleotide STRs are the most common categories of STRs in the vertebrate genomes[20, 21]. In addition to their association with frameshifts in coding sequences and pathological [22] and possibly evolutionary consequences, recent evidence indicates surprising functions for the mononucleotide STRs, such as their provisional role in translation initiation site selection[9]. Several groups have found evidence on the involvement of a number of dinucleotide STRs in gene regulation, speciation, and evolution[3, 20, 23–26]. Trinucleotide STRs are frequently linked to human neurological disorders, most of which are specific to this species[27, 28].

In a model study, here we analyzed the evolutionary abundance of all types of mono-, di-, and trinucleotide STRs in nine selected species, encompassing rodents, Old World monkeys, and great apes.

## Materials and Methods

### Species and whole-genome sequences

By using the UCSC genome browser (https://hgdownload.soe.ucsc.edu), the whole genomes of nine species were downloaded and analyzed, species and genome sizes of which were as follows: rat (*Rattus norvegicus*): 2,647,915,728, mouse (*Mus musculus*): 2,728,222,451, gelada (*Theropithecus gelada*): 2,889,630,685, olive baboon (*Papio anubis*): 2,869,821,163, macaque (*Macaca mulatta*): 2,946,843,737, gorilla (*Gorilla gorilla gorilla*): 3,063,362,754, chimpanzee (*Pan troglodytes*): 3,050,398,082, bonobo (*Pan paniscus*): 3,203,531,224, and human (*Homo sapiens*): 3,099,706,404. Those species encompassed rodents: rat and mouse, Old World monkeys: gelada, olive baboon, macaque, and great apes: gorilla, bonobo, chimpanzee, human.

### Extraction of STRs from genomic sequences

The whole-genome abundance of mononucleotide STRs of ≥10-repeats, dinucleotide STRs of ≥6-repeats, and trinucleotide STRs of ≥4-repeats were studied in the nine selected species. To that end, we designed a software package in Java (https://github.com/arabfard/Java_STR_Finder). All possibilities of mononucleotide motifs, consisting of A, C, T, and G, all possibilities of dinucleotide motifs, consisting of AC, AG, AT, CA, CG, CT, GA, GC, GT, TA, TC, and TG, and all possibilities of trinucleotide motifs, consisting of AAC, AAT, AAG, ACA, ACC, ACT, ACG, ATA, ATC, ATT, ATG, AGA, AGC, AGT, AGG, CAA, CAC, CAT, CAG, CCA, CCT, CCG, CTA, CTC, CTT, CTG, CGA, CGC, CGT, CGG, TAA, TAC, TAT, TAG, TCA, TCC, TCT, TCG, TTA, TTC, TTG, TGA, TGC, TGT, TGG, GAA, GAC, GAT, GAG, GCA, GCC, GCT, GCG, GTA, GTC, GTT, GTG, GGA, GGC, and GGT were analyzed. The written program was based on perfect (pure) STRs.

### Chromosome-by-chromosome aggregation of STRs

Whole-genome chromosome-by-chromosome data were aggregated and analyzed in the nine species, without normalization (approach 1) and with normalization (approach 2). In approach 1, all chromosomal data were collected without removing any numerically non-identical chromosomes across the nine species. In approach 2, data on the identical chromosome sets (numerically) across the nine species were collected in an array of 20 columns, each column corresponding to a chromosome. In this approach, mouse was selected as reference, because it had the lowest number of chromosomes among the nine species.

### STR abundance and hierarchical cluster analysis across species

Whole-genome STR abundances across the selected species were deciphered and depicted by boxplot diagrams and hierarchical clustering, using boxplot and hclust packages[29] in R, respectively. Boxplots illustrate abundance differences among segments across the selected species, and hierarchical clustering plots demonstrate the level of similarity and differences across the obtained abundances. The input data to these packages were numerical arrays obtained with each approach. Each array consisted of a number of columns, each column corresponding to the STR abundance in different chromosomes.

### Statistical analysis

The STR abundances across the nine selected species were compared by repeated measurements analysis, using one and two-way ANOVA tests. These analyses were confirmed by nonparametric tests.

## Results

### Global abundance of mono, di, and trinucleotide STRs coincides with the phylogenetic distance of the nine selected species

Chromosome-by-chromosome data were collected on the abundance of mononucleotide STRs across the nine species (Table 1). We found massive expansion of the mononucleotide STR compartment in all primate species vs. rat and mouse. Hierarchical clustering yielded three <clusters> as follows: <rat, mouse>, <gelada, olive baboon, macaque>, and <gorilla, chimpanzee, bonobo, human>, which coincided with the phylogenetic distance of the nine selected species in both unnormalized (P=6.3E-09) (Fig. 1) and normalized approaches (P=1.4E-08) (Suppl. 1), namely <rodents>, <Old World monkeys>, and <great apes>.

**Fig. 1.**
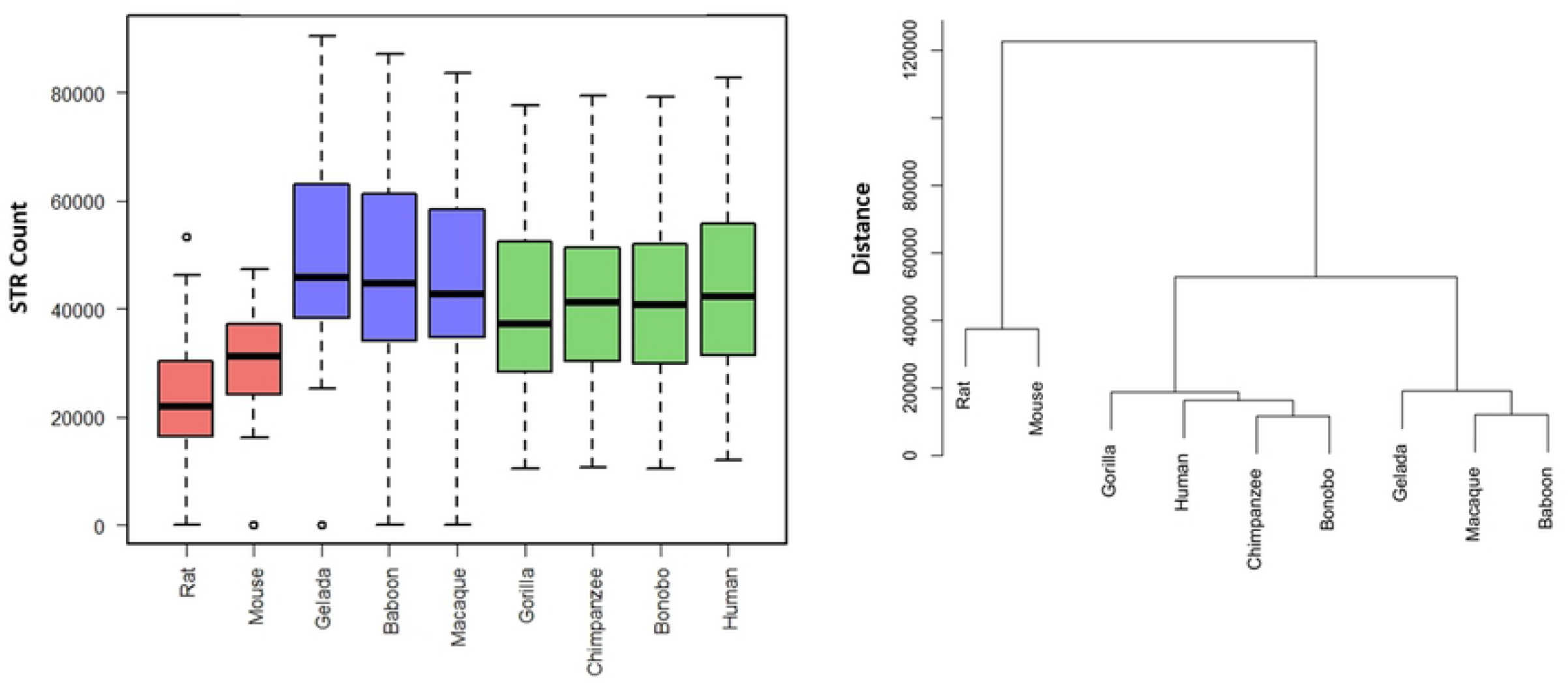
Unnormalized data on whole-genome mononucleotide STRs in the nine selected species. Global incremented pattern was observed in the primate species vs. rodents (left graphs). The overall hierarchical clustering yielded three clusters, which coincided with rodents, Old World monkeys, and great apes (right graphs).

**Table 1.** Whole-genome mononucleotide STR abundance. Chromosome-by-chromosome data across the nine selected species.

| Chromosome/Species | Rat | Mouse | Gelada | Baboon | Macaque | Gorilla | Chimpanzee | Bonobo | Human |
| --- | --- | --- | --- | --- | --- | --- | --- | --- | --- |
| **1** | 53318 | 47294 | 90549 | 87241 | 83595 | 77718 | 79390 | 79173 | 82820 |
| **2(A)** | 46221 | 45636 | 71588 | 67963 | 64609 | 35908 | 35897 | 34400 | 78550 |
| **2(B)** | 0 | 0 | 0 | 0 | 0 | 40245 | 39968 | 39837 | 0 |
| **3** | 36364 | 38493 | 70736 | 68688 | 65836 | 62398 | 62713 | 64472 | 64027 |
| **4** | 34818 | 39019 | 62831 | 60726 | 57817 | 54896 | 54855 | 53287 | 56495 |
| **5** | 36532 | 38805 | 66164 | 64101 | 61533 | 60436 | 48944 | 54142 | 56538 |
| **6** | 28617 | 35751 | 63104 | 61642 | 59150 | 53872 | 53769 | 53420 | 55185 |
| **7(A)** | 29411 | 33649 | 25699 | 65267 | 63438 | 50898 | 53882 | 50792 | 56257 |
| **7(B)** | 0 | 0 | 42663 | 0 | 0 | 0 | 0 | 0 | 0 |
| **8** | 27353 | 31938 | 50576 | 48446 | 46757 | 43593 | 44212 | 43618 | 45220 |
| **9** | 23532 | 31142 | 50050 | 47879 | 46910 | 36797 | 38035 | 37493 | 41744 |
| **10** | 31065 | 34138 | 41475 | 39012 | 37477 | 44166 | 44562 | 44416 | 46075 |
| **11** | 17071 | 33869 | 54287 | 54284 | 51654 | 37218 | 41059 | 40757 | 42217 |
| **12** | 15101 | 29325 | 42675 | 35365 | 42793 | 46865 | 47576 | 47481 | 48483 |
| **13** | 21673 | 29496 | 40602 | 39101 | 38022 | 27902 | 28481 | 28479 | 29430 |
| **14** | 21835 | 28835 | 45820 | 44693 | 42677 | 30311 | 30659 | 30595 | 31460 |
| **15** | 20351 | 25753 | 43334 | 41671 | 40009 | 28611 | 29752 | 29049 | 31402 |
| **16** | 15958 | 24139 | 41211 | 39781 | 37693 | 29268 | 31121 | 28460 | 34364 |
| **17** | 18458 | 24234 | 32308 | 31285 | 30378 | 29884 | 36791 | 37010 | 38947 |
| **18** | 16651 | 22580 | 25310 | 24850 | 23551 | 22556 | 22428 | 22236 | 23130 |
| **19** | 14266 | 16221 | 35819 | 32702 | 30470 | 23832 | 31405 | 30614 | 32423 |
| **20** | 14475 | 0 | 34962 | 32965 | 32095 | 20654 | 22106 | 31034 | 21961 |
| **21** | 0 | 0 | 0 | 0 | 0 | 10462 | 10633 | 10467 | 12050 |
| **22** | 0 | 0 | 0 | 0 | 0 | 13778 | 14816 | 13904 | 16014 |
| **X** | 25983 | 40547 | 52836 | 49013 | 47590 | 43138 | 43302 | 41656 | 46178 |
| **Sum** | **549053** | **650864** | **1084599** | **1036675** | **1004054** | **925406** | **946356** | **946792** | **990970** |

The whole-genome STR abundances from aggregated chromosome-by-chromosome analysis in the dinucleotide category (Table 2) was decremented in primates vs. rodents. Similar to the mononucleotide STR compartment, the dinucleotide STR compartment coincided with the genetic distance among the three <clusters> of species with the unnormalized (P=7.1E-08) (Fig. 2) and normalized data (P=6.8E-11) (Suppl. 1).

**Fig. 2.**
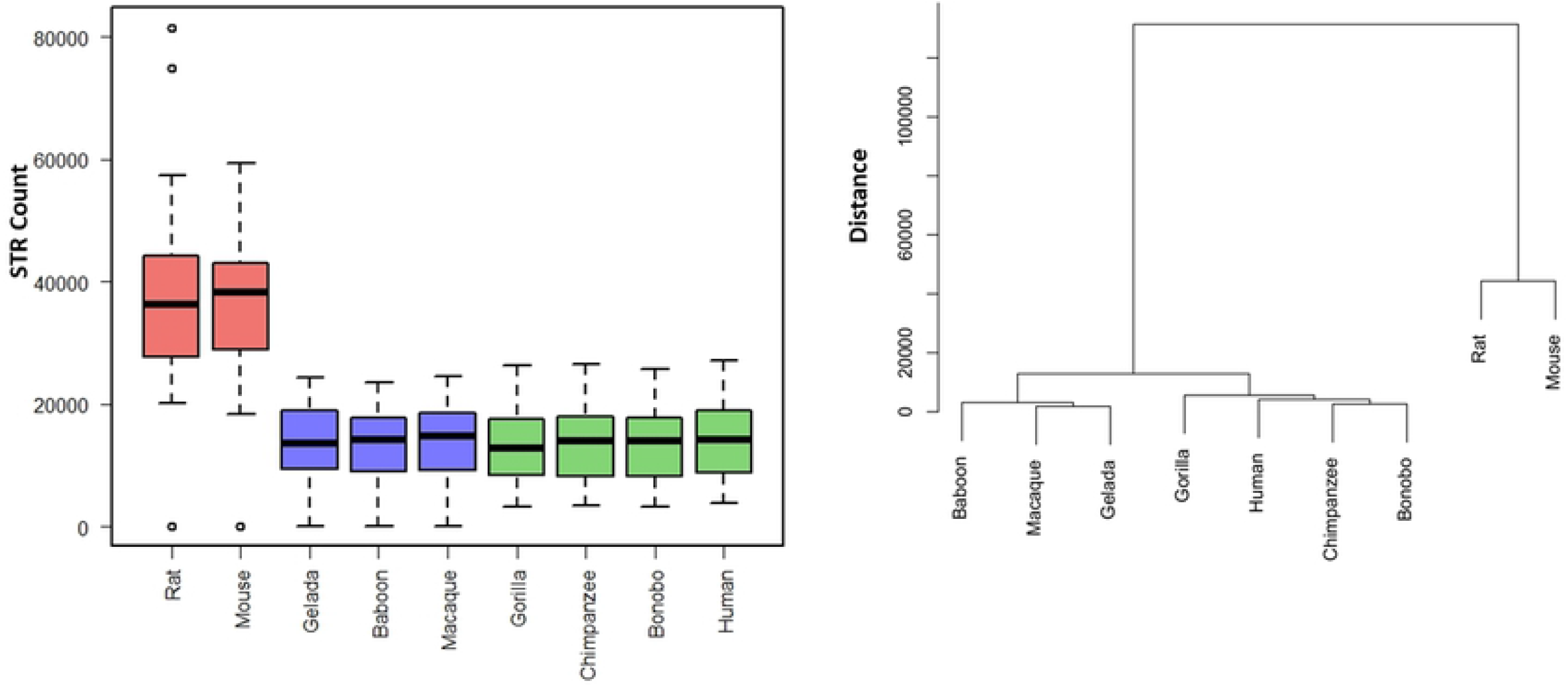
Unnormalized data on whole-genome dinucleotide STRs in the nine selected species. Global decremented patterns were observed in all primate species vs. mouse and rat.

**Table 2.** Whole-genome dinucleotide STR abundance. Chromosome-by-chromosome data across the nine selected species.

| Chromosome/Species | Rat | Mouse | Gelada | Baboon | Macaque | Gorilla | Chimpanzee | Bonobo | Human |
| --- | --- | --- | --- | --- | --- | --- | --- | --- | --- |
| **1** | 81509 | 59425 | 24335 | 23427 | 24462 | 23105 | 23708 | 23583 | 24657 |
| **2(A)** | 74837 | 53096 | 21315 | 20302 | 21225 | 11820 | 11960 | 11391 | 26989 |
| **2(B)** | 0 | 0 | 0 | 0 | 0 | 14494 | 14555 | 14334 | 0 |
| **3** | 53642 | 45464 | 20710 | 19973 | 20552 | 20939 | 21179 | 21039 | 21633 |
| **4** | 57299 | 44963 | 19364 | 18592 | 19038 | 21536 | 21182 | 20503 | 21773 |
| **5** | 52269 | 48069 | 22020 | 21275 | 22147 | 17099 | 17831 | 19606 | 20385 |
| **6** | 44993 | 45325 | 19921 | 19397 | 20070 | 18575 | 18391 | 18196 | 18995 |
| **7(A)** | 43219 | 40052 | 5832 | 16963 | 17870 | 15988 | 16727 | 16130 | 17275 |
| **7(B)** | 0 | 0 | 11934 | 0 | 0 | 0 | 0 | 0 | 0 |
| **8** | 43242 | 41103 | 15903 | 15390 | 16164 | 15837 | 15875 | 15718 | 16245 |
| **9** | 37463 | 39005 | 14733 | 14183 | 14857 | 11704 | 11935 | 11661 | 13080 |
| **10** | 40260 | 40998 | 10136 | 9432 | 9855 | 14051 | 14306 | 14032 | 14799 |
| **11** | 27685 | 38212 | 14360 | 14487 | 15187 | 12678 | 13988 | 13842 | 14189 |
| **12** | 22084 | 35361 | 13478 | 14325 | 14685 | 14385 | 14559 | 14588 | 14757 |
| **13** | 38331 | 35159 | 11839 | 11292 | 11797 | 11071 | 11258 | 11135 | 11406 |
| **14** | 31923 | 36644 | 13605 | 13243 | 13885 | 9549 | 9465 | 9386 | 9798 |
| **15** | 31768 | 30662 | 12078 | 11661 | 12014 | 8014 | 8226 | 8143 | 8607 |
| **16** | 28704 | 29521 | 8228 | 8064 | 8206 | 7814 | 8268 | 7553 | 8947 |
| **17** | 30312 | 28209 | 11002 | 10457 | 10942 | 10456 | 8056 | 8006 | 8355 |
| **18** | 27797 | 27263 | 8548 | 8349 | 8591 | 8629 | 8597 | 8497 | 8750 |
| **19** | 21794 | 18350 | 5994 | 5493 | 5395 | 4774 | 6081 | 5865 | 6220 |
| **20** | 20191 | 0 | 8334 | 7902 | 8345 | 6379 | 7106 | 6623 | 6612 |
| **21** | 0 | 0 | 0 | 0 | 0 | 4092 | 4154 | 4123 | 4884 |
| **22** | 0 | 0 | 0 | 0 | 0 | 3209 | 3442 | 3183 | 3746 |
| **X** | 36246 | 38470 | 18303 | 16787 | 17659 | 17922 | 18193 | 17078 | 18952 |
| **Sum** | **845568** | **775351** | **311972** | **300994** | **312946** | **304120** | **309042** | **304215** | **321054** |

There was global shrinkage of the trinucleotide STR compartment in primates vs. rodents, without (P=3.8E-05) and with normalization of the data (P=2.4E-07) (Table 3, Fig. 3 and Suppl. 1). Remarkably, human stood out among all other species in the hierarchical clustering,

**Fig. 3.**
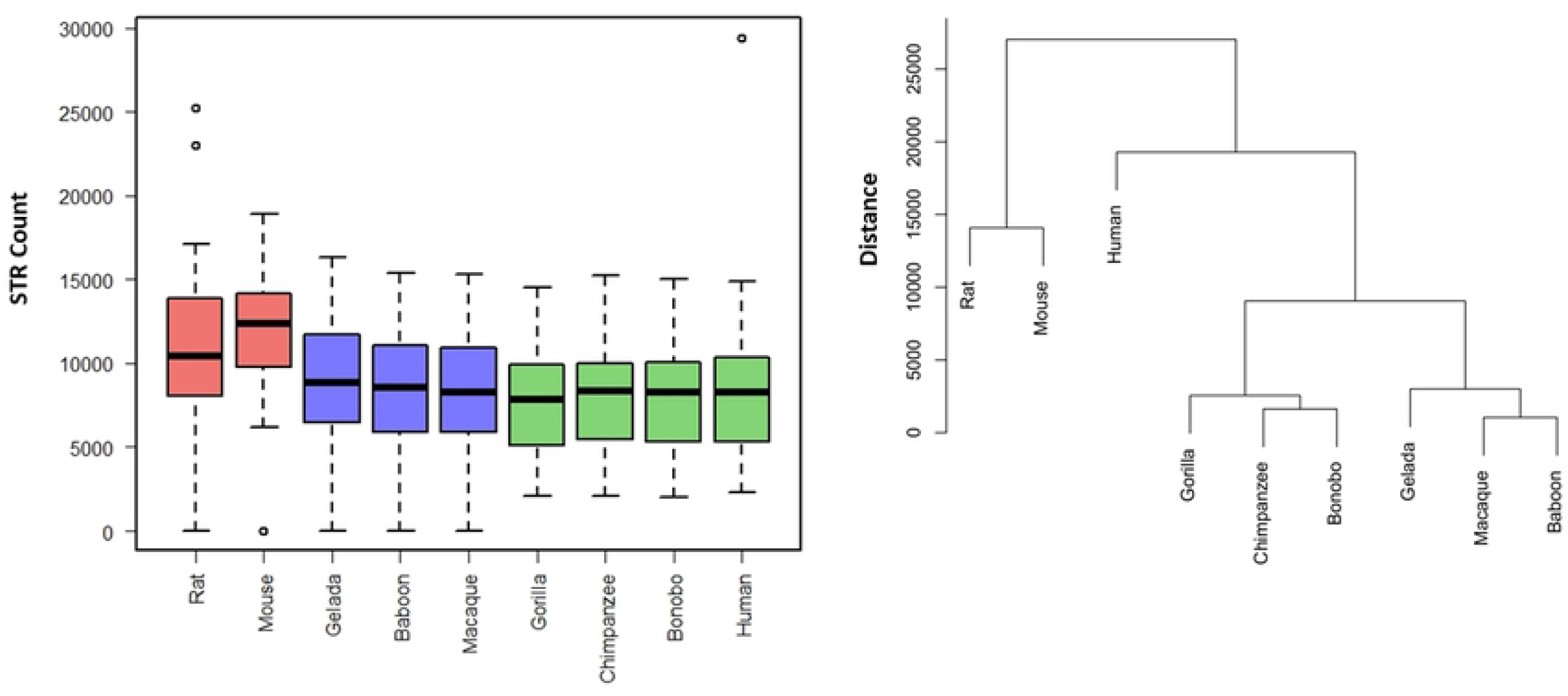
Unnormalized data on whole-genome trinucleotide STRs in the nine selected species. While global decremented patterns were observed in primates vs. rodents, intriguingly, human stood out in this category, in comparison to all other species.

**Table 3.** Whole-genome trinucleotide STR abundance. Chromosome-by-chromosome data across the nine selected species.

| Chromosome/Species | Rat | Mouse | Gelada | Baboon | Macaque | Gorilla | Chimpanzee | Bonobo | Human |
| --- | --- | --- | --- | --- | --- | --- | --- | --- | --- |
| **1** | 25234 | 18913 | 16307 | 15350 | 15341 | 14540 | 15219 | 15054 | 14882 |
| **2(A)** | 22996 | 17856 | 13005 | 12341 | 11998 | 6800 | 6842 | 6537 | 14521 |
| **2(B)** | 0 | 0 | 0 | 0 | 0 | 7545 | 7764 | 7822 | 0 |
| **3** | 16869 | 15022 | 12749 | 12518 | 11938 | 11473 | 11744 | 11637 | 11631 |
| **4** | 17088 | 15204 | 11921 | 11154 | 10960 | 11116 | 11228 | 10685 | 11144 |
| **5** | 16339 | 15469 | 13001 | 12514 | 12112 | 10581 | 9665 | 10640 | 10649 |
| **6** | 13495 | 14332 | 12150 | 11743 | 11380 | 10364 | 10504 | 10445 | 29430 |
| **7(A)** | 14317 | 13760 | 3937 | 10991 | 10871 | 9342 | 10117 | 9744 | 9995 |
| **7(B)** | 0 | 0 | 7552 | 0 | 0 | 0 | 0 | 0 | 0 |
| **8** | 12701 | 13518 | 10032 | 9524 | 9682 | 8752 | 9096 | 8645 | 8890 |
| **9** | 11646 | 12378 | 9295 | 8755 | 8659 | 6898 | 7328 | 7157 | 7580 |
| **10** | 12552 | 13968 | 7297 | 6728 | 6786 | 8096 | 8350 | 8245 | 8295 |
| **11** | 7987 | 13232 | 9615 | 9578 | 9403 | 7801 | 8668 | 8458 | 8352 |
| **12** | 6060 | 11817 | 7742 | 8297 | 8029 | 8905 | 9218 | 9051 | 9127 |
| **13** | 10852 | 11634 | 7266 | 6823 | 6860 | 5273 | 5479 | 5452 | 5391 |
| **14** | 10325 | 11865 | 8869 | 8583 | 8253 | 5473 | 5771 | 5785 | 5706 |
| **15** | 10075 | 10693 | 7727 | 7339 | 7152 | 4869 | 5168 | 5082 | 5297 |
| **16** | 8476 | 9527 | 6228 | 5837 | 5801 | 5738 | 6007 | 5623 | 6402 |
| **17** | 9502 | 10045 | 5908 | 5737 | 5684 | 5666 | 5859 | 5914 | 6091 |
| **18** | 8124 | 9154 | 4738 | 4645 | 4603 | 4722 | 4625 | 4584 | 4566 |
| **19** | 6984 | 6190 | 5432 | 4643 | 4664 | 3807 | 5438 | 5230 | 5101 |
| **20** | 6445 | 0 | 6655 | 6016 | 5945 | 4072 | 4472 | 4155 | 4130 |
| **21** | 0 | 0 | 0 | 0 | 0 | 2051 | 2092 | 2028 | 2304 |
| **22** | 0 | 0 | 0 | 0 | 0 | 2721 | 2825 | 2601 | 2915 |
| **X** | 10411 | 13783 | 11449 | 10609 | 10666 | 9547 | 9838 | 9140 | 10062 |
| **Sum** | **258478** | **258360** | **198875** | **189725** | **186787** | **176152** | **183317** | **179714** | **202461** |

### Differential abundance patterns of STRs across rodents and primates

Numerous STRs across the mono, di, and trinucleotide STR categories coincided with the phylogenetic distances of the nine selected species. For example, the most abundant STRs across all nine species were T/A mononucleotides of 10, 11, and 12 repeats, which coincided with the genetic distance of the selected species (Fig. 4). Likewise, (ct)6 and (taa)4 conformed to the phylogeny of the studied species in the di and trinucleotide STR categories, respectively. On the other hand, numerous STRs did not follow a phylogenetic pattern, such as (C)10, (AT)8, and (ttg)4 (Fig. 5). Hierarchical clusters of all studied STRs across the three categories are available at: https://figshare.com/articles/figure/STR_Clustering/17054972.

**Fig. 4.**
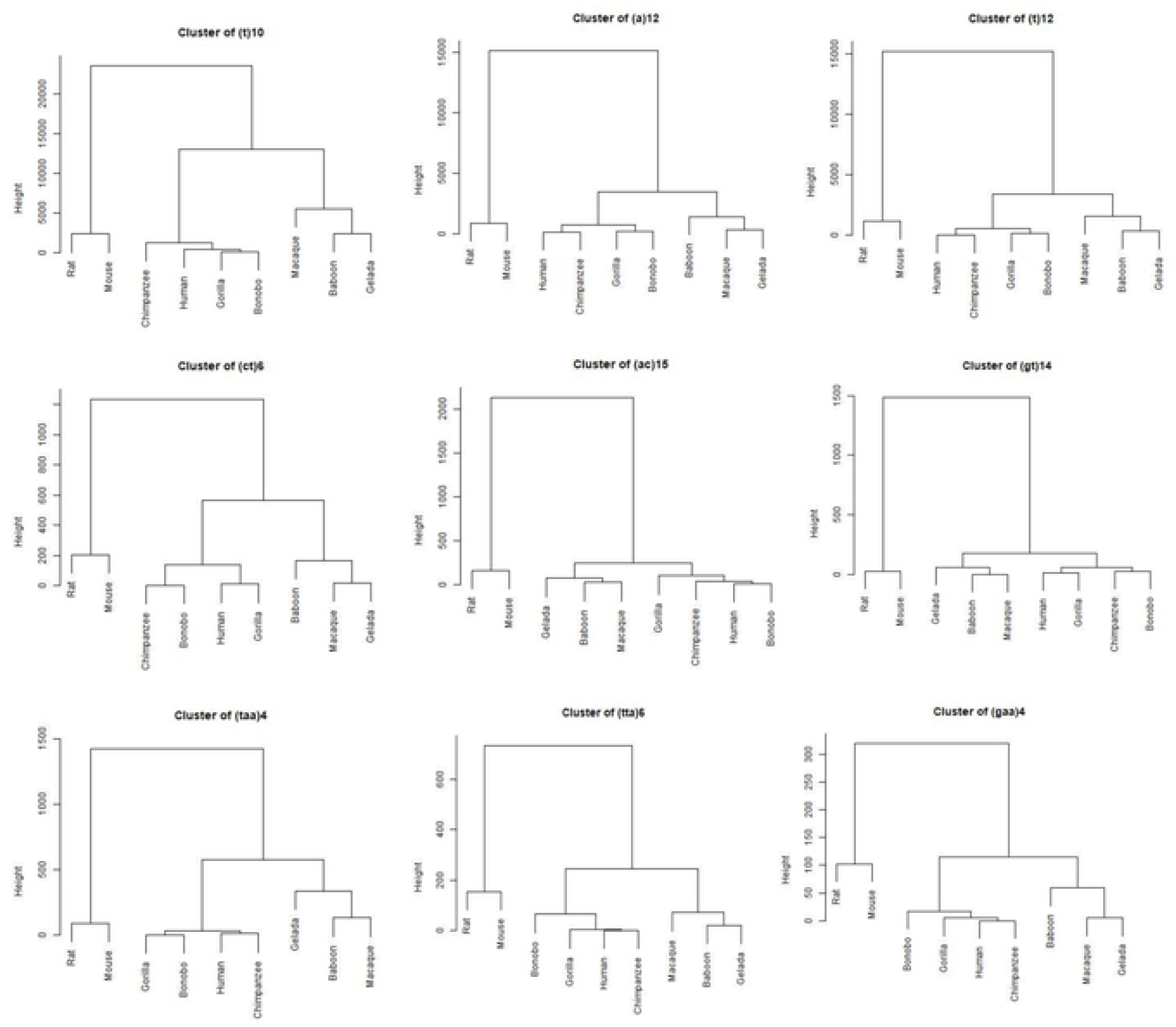
Example of STRs and STR lengths, abundance of which coincided with the phylogeny of the nine selected species. Three STRs are depicted as examples for each of mono, di, and trinucleotide categories. Data from all studied STRs are available at: https://figshare.com/articles/figure/STR_Clustering/17054972.

**Fig. 5.**
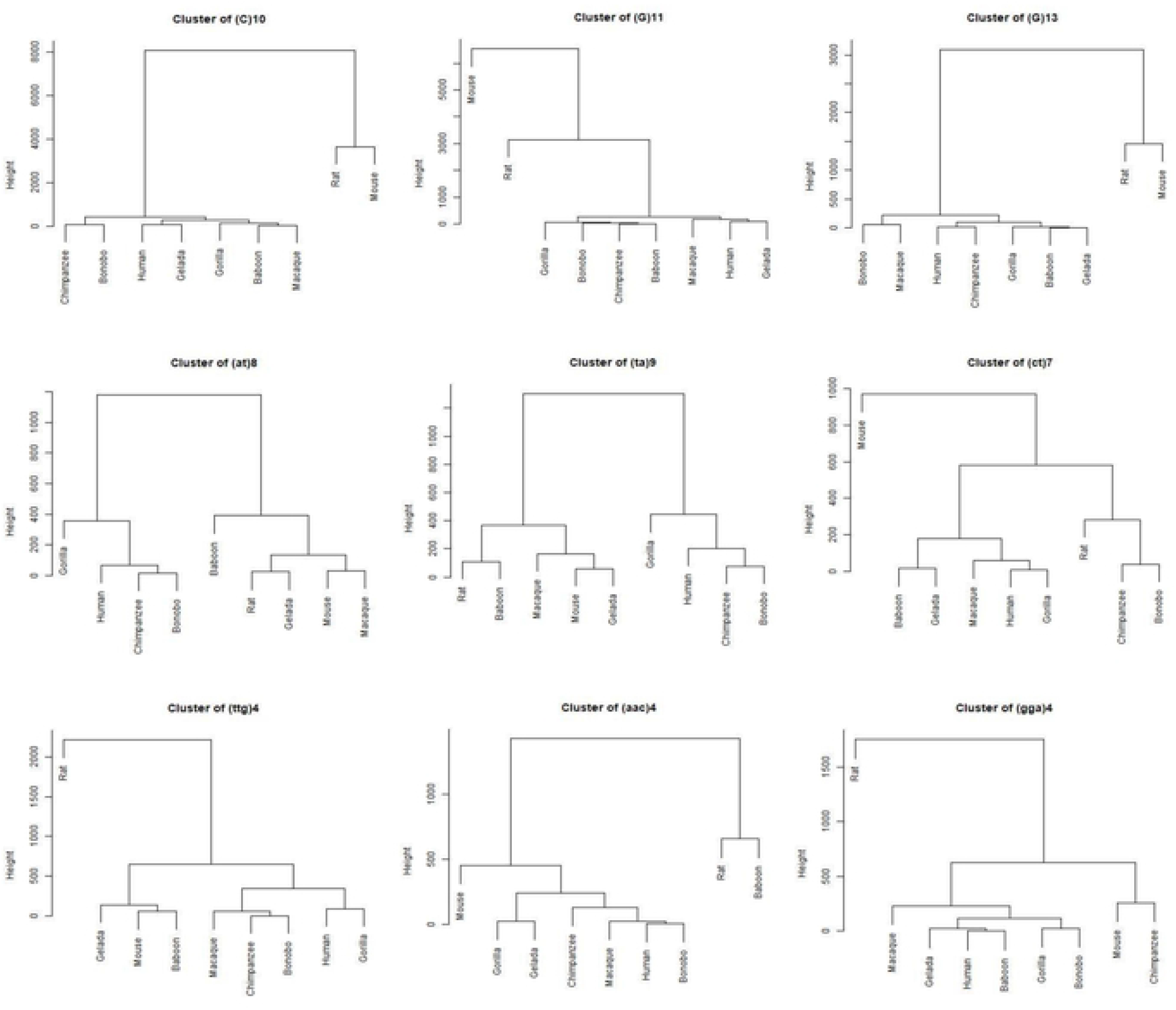
Example of STRs and STR lengths, abundance of which appeared to be random across the nine selected species. Three STRs are depicted as examples for each of mono, di, and trinucleotide categories. Data from all studied STRs are available at: https://figshare.com/articles/figure/STR_Clustering/17054972.

## Discussion

It is largely unknown whether at the crossroads of speciation, STRs evolved as a result of purifying selection, genetic drift, and/or in a directional manner. In a model study, we selected multiple species across rodents and primates, and investigated the abundance of all possible types of mononucleotides, dinucleotide, and trinucleotide STRs on the whole-genome scale in those species. Hierarchical clustering of the obtained abundances yielded clusters that predominantly coincided with the phylogenetic distances of the selected species.

Hierarchical clustering is an unsupervised clustering method that is used to group data. This algorithm is unsupervised because it uses random, unlabelled datasets. As the number of clusters increases, the accuracy of the hierarchical clustering algorithm improves. Here we implemented this algorithm to cluster the nine selected species based on the obtained STR abundances. Our findings may be of significance in two respects. Firstly, there were significant differential abundances separating rodents from primates, for example, massive decremented abundance of dinucleotide and trinucleotide STRs in primates vs. the rodent species, and massive incremented abundance of mononucleotide STRs in primates vs. rodents. Those differential abundances might have determining roles in the speciation of the two orders. Secondly, the three major clusters obtained from global hierarchical cluster analysis matched the phylogeny of the three classes of species, i.e., <rodents>, <Old World monkeys>, and <great apes>. It is possible that there are mathematical channels/thresholds required for the abundance of STRs in various orders. This is in line with the hypothesis that STRs function as scaffolds for biological computers[30].

In addition, our data indicate that various STRs and STR lengths behave differently with respect to their colossal abundance. Not all studied STRs coincided with the phylogenetic distances of the nine selected species. We hypothesize that those which coincided had a link with the speciation of those species, whereas those which did not probably followed random patterns. The obtained abundances seem to be independent of the genome sizes of the selected species, for example in the instances of di- and trinucleotide STRs. This finding is in line with the previous reports of lack of relationship between genome size and abundance of STRs[14, 31, 32]. Mononucleotide STRs impact various processes, such as gene expression and translation alterations and frameshifts of various proteins, which may have evolutionary and pathological consequences[9, 22]. They can overlap with G4 structures, many of which associate with evolutionary consequences[33].

Dinucleotide STRs located in the protein-coding gene core promoters have been subject to contraction in a number of instances, in the process of human and non-human primate evolution[34]. A number of those STRs are identical in formula in primates vs. non-primates, and the genes linked to those STRs are involved in characteristics that have diverged primates from other mammals, such as craniofacial development, neurogenesis, and spine morphogenesis. It is likely that those STRs functioned as evolutionary switch codes for primate speciation. In line with the above, structural variants are enriched near genes that diverged in expression across great apes[35], and genes with STRs in their regulatory regions were more divergent in expression than genes with fixed or no STRs[36]. It is speculated that STR variants are more likely than single-nucleotide variants to have epistatic interactions, which can have significant consequences in complex traits, in human as well as model organisms[37, 38]. Trinucleotide STRs are predominantly focused on in human because of their link with several neurological disorders[39–44]. Intriguingly, we found an exceptional global hierarchical distance between human and all other species. In view of the fact that most of the phenotypes attributed to trinucleotide STRs are human-specific in nature, it is conceivable that their evolution is also significantly distant from all other species studied.

Future studies such as large-scale genome-editing of STRs[45] in embryonic stem cells and investigation of their differentiation into various cell lineages may be candidate approaches to investigate how the observed massive non-random patterns link to speciation and evolution.

## Conclusion

We propose that the global abundance of STRs is non-random across rodents and primates. We also propose the STRs and STR lengths which coincided with the phylogenetic distances of those species.

## Declarations

### Ethics approval and consent to participate

Not applicable

### Consent for publication

Not applicable

### Availability of data and materials

Raw data are available at: https://figshare.com/articles/dataset/Trends/15073329 and https://figshare.com/articles/figure/STR_Clustering/17054972

### Authors’ contributions

MA performed and coordinated the bioinformatics analyses. MS performed the biostatistics analysis. YHN, IA, and AMAM contributed to data collection. KK contributed to data collection and coordination. MO conceived and supervised the project, and wrote the manuscript.

## Acknowledgements

Not applicable.

## Figures

**Suppl. 1.** Normalized data across the nine selected species.

## References

1. Mohammadparast, S., et al., Exceptional expansion and conservation of a CT-repeat complex in the core promoter of PAXBP1 in primates. American journal of primatology, 2014. 76(8): p. 747–756.

2. Bushehri, A., et al., Genome-wide identification of human-and primate-specific core promoter short tandem repeats. Gene, 2016. 587(1): p. 83–90.

3. Nikkhah, M., et al., An exceptionally long CA-repeat in the core promoter of SCGB2B2 links with the evolution of apes and Old World monkeys. Gene, 2016. 576(1): p. 109–114.

4. Jakubosky, D., et al., Properties of structural variants and short tandem repeats associated with gene expression and complex traits. Nature Communications, 2020. 11(1): p. 1–15.

5. Valipour, E., et al., Polymorphic core promoter GA-repeats alter gene expression of the early embryonic developmental genes. Gene, 2013. 531(2): p. 175–179.

6. Ranathunge, C., et al., Transcribed microsatellite allele lengths are often correlated with gene expression in natural sunflower populations. Molecular Ecology, 2020.

7. Press, M.O., et al., Substitutions are boring: Some arguments about parallel mutations and high mutation rates. Trends in Genetics, 2019. 35(4): p. 253–264.

8. Fotsing, S.F., et al., The impact of short tandem repeat variation on gene expression. Nature genetics, 2019. 51(11): p. 1652–1659.

9. Arabfard, M., et al., Link between short tandem repeats and translation initiation site selection. Human genomics, 2018. 12(1): p. 47.

10. Yap, K., et al., A short tandem repeat-enriched RNA assembles a nuclear compartment to control alternative splicing and promote cell survival. Molecular cell, 2018. 72(3): p. 525–540.

11. Fondon, J.W. and H.R. Garner, Molecular origins of rapid and continuous morphological evolution. Proceedings of the National Academy of Sciences, 2004. 101(52): p. 18058–18063.

12. Wren, J.D., et al., Repeat polymorphisms within gene regions: phenotypic and evolutionary implications. The American Journal of Human Genetics, 2000. 67(2): p. 345–356.

13. King, D.G., Evolution of simple sequence repeats as mutable sites. Tandem Repeat Polymorphisms, 2012: p. 10–25.

14. Srivastava, S., et al., Patterns of microsatellite distribution across eukaryotic genomes. BMC genomics, 2019. 20(1): p. 153.

15. Pavlova, A., et al., Purifying selection and genetic drift shaped Pleistocene evolution of the mitochondrial genome in an endangered Australian freshwater fish. Heredity, 2017. 118(5): p. 466–476.

16. Jorde, P.E., et al., Genetically distinct populations of northern shrimp, Pandalus borealis, in the North Atlantic: adaptation to different temperatures as an isolation factor. Molecular Ecology, 2015. 24(8): p. 1742–1757.

17. Legrand, D., et al., Inter-island divergence within Drosophila mauritiana, a species of the D. simulans complex: Past history and/or speciation in progress? Molecular Ecology, 2011. 20(13): p. 2787–2804.

18. Sun, G., et al., Global genetic variation at nine short tandem repeat loci and implications on forensic genetics. European Journal of Human Genetics, 2003. 11(1): p. 39–49.

19. Abe, H. and N.J. Gemmell, Evolutionary footprints of short tandem repeats in avian promoters. Scientific reports, 2016. 6(1): p. 1–11.

20. Lander, E.S., et al., Initial sequencing and analysis of the human genome. 2001.

21. Fan, H. and J.-Y. Chu, A brief review of short tandem repeat mutation. Genomics, proteomics & bioinformatics, 2007. 5(1): p. 7–14.

22. Mo, H.Y., et al., Frameshift Mutations and Loss of Expression of CLCA4 Gene are Frequent in Colorectal Cancers With Microsatellite Instability. Applied Immunohistochemistry & Molecular Morphology, 2020. 28(7): p. 489.

23. Corney, B.P.A., et al., Regulatory architecture of the neuronal Cacng2/Tarpγ2 gene promoter: multiple repressive domains, a polymorphic regulatory short tandem repeat, and bidirectional organization with co-regulated lncRNAs. Journal of Molecular Neuroscience, 2019. 67(2): p. 282–294.

24. Emamalizadeh, B., et al., The human RIT2 core promoter short tandem repeat predominant allele is species-specific in length: a selective advantage for human evolution? Molecular Genetics and Genomics, 2017. 292(3): p. 611–617.

25. Haasl, R.J., R.C. Johnson, and B.A. Payseur, The effects of microsatellite selection on linked sequence diversity. Genome biology and evolution, 2014. 6(7): p. 1843–1861.

26. Yim, J.-J., et al., Evolution of an intronic microsatellite polymorphism in Toll-like receptor 2 among primates. Immunogenetics, 2006. 58(9): p. 740–745.

27. Annear, D.J., et al., Abundancy of polymorphic CGG repeats in the human genome suggest a broad involvement in neurological disease. Scientific reports, 2021. 11(1): p. 1–11.

28. Tang, H., et al., Profiling of short-tandem-repeat disease alleles in 12,632 human whole genomes. The American Journal of Human Genetics, 2017. 101(5): p. 700–715.

29. Murtagh, F. and P. Legendre, Ward’s hierarchical agglomerative clustering method: which algorithms implement Ward’s criterion? Journal of classification, 2014. 31(3): p. 274–295.

30. Herbert, A., Simple Repeats as Building Blocks for Genetic Computers. Trends in Genetics, 2020.

31. Neff, B.D. and M.R. Gross, Microsatellite evolution in vertebrates: inference from AC dinucleotide repeats. Evolution, 2001. 55(9): p. 1717–1733.

32. Park, J.Y., et al., Evolutionary constraints over microsatellite abundance in larger mammals as a potential mechanism against carcinogenic burden. Scientific reports, 2016. 6(1): p. 1–5.

33. Sawaya, S., et al., Microsatellite tandem repeats are abundant in human promoters and are associated with regulatory elements. PloS one, 2013. 8(2): p. e54710.

34. Ohadi, M., et al., Core promoter short tandem repeats as evolutionary switch codes for primate speciation. American journal of primatology, 2015. 77(1): p. 34–43.

35. Kronenberg, Z.N., et al., High-resolution comparative analysis of great ape genomes. Science, 2018. 360(6393).

36. Sonay, T.B., et al., Tandem repeat variation in human and great ape populations and its impact on gene expression divergence. Genome research, 2015. 25(11): p. 1591–1599.

37. Bagshaw, A.T.M., et al., Microsatellite polymorphisms associated with human behavioural and psychological phenotypes including a gene-environment interaction. BMC medical genetics, 2017. 18(1): p. 1–12.

38. Press, M.O., K.D. Carlson, and C. Queitsch, The overdue promise of short tandem repeat variation for heritability. Trends in Genetics, 2014. 30(11): p. 504–512.

39. Khamse S, Jafarian Z, Bozorgmehr A, Tavakoli M, Afshar H, Keshavarz M, Moayedi R, Ohadi M. Novel implications of a strictly monomorphic (GCC) repeat in the human PRKACB gene. Sci Rep. 2021 Oct 19;11(1):20629. doi: 10.1038/s41598-021-99932-3. PMID: 34667254; PMCID: PMC8526596.

40. Jafarian Z, Khamse S, Afshar H, Khorshid HRK, Delbari A, Ohadi M. Natural selection at the RASGEF1C (GGC) repeat in human and divergent genotypes in late-onset neurocognitive disorder. Sci Rep. 2021 Sep 28;11(1):19235. doi: 10.1038/s41598-021-98725-y. PMID: 34584172; PMCID: PMC8479062.

41. Sundblom, J., et al., High frequency of intermediary alleles in the HTT gene in Northern Sweden-The Swedish Huntingtin Alleles and Phenotype (SHAPE) study. Scientific reports, 2020. 10(1): p. 1–7.

42. Baker, E.K., et al., FMR1 mRNA from full mutation alleles is associated with ABC-C FX scores in males with fragile X syndrome. Scientific Reports, 2020. 10(1): p. 1–8.

43. Zhou, X., et al., Analysis of (CAG) n expansion in ATXN1, ATXN2 and ATXN3 in Chinese patients with multiple system atrophy. Scientific reports, 2018. 8(1): p. 1–5.

44. Zhang, Q., et al., A brain-targeting lipidated peptide for neutralizing RNA-mediated toxicity in Polyglutamine Diseases. Scientific reports, 2017. 7(1): p. 1–13.

45. Smith, C.J., et al., Enabling large-scale genome editing at repetitive elements by reducing DNA nicking. Nucleic acids research, 2020. 48(9): p. 5183–5195.

